# Statistical and structural bias in birth-death models

**DOI:** 10.64898/2025.12.02.691894

**Authors:** Jeremy M. Beaulieu, Brian C. O’Meara

## Abstract

Accurate estimation of speciation (***λ***) and extinction (***µ***) rates from phylogenetic trees is central to studies of diversification, yet it remains unclear whether commonly used estimators are unbiased. Here we examine two sources of error: (1) statistical bias in the estimators themselves, and the (2) structural bias introduced by how small trees are handled in likelihood calculations. For the Yule process, we re-derive the expected bias of the standard estimator, showing that 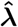 underestimates ***λ*** by a factor of **(*n* − 2)/(*n* − 1)**. Extending to the general birth–death model, we use symbolic regression to find functional forms that minimize the bias in both ***λ*** and ***µ***. The best-performing correction for ***λ*** is identical to the Yule result, while the bias in ***µ*** depends on both sample size and the estimated extinction fraction (***µ/λ***). Applying these corrections substantially improves the fit between the estimated and generating values. When these corrected estimators are used to derive other diversification-related parameters, turnover is nearly unbiased, but net diversification (***λ* − *µ***) remains systematically underestimated due to the slight overestimation of ***µ***. On the whole, these results clarify the statistical and structural sources of bias in diversification rate estimation and provide a general framework for improving inference under birth-death models.

## 1 Introduction

Understanding and uncovering the dynamics of speciation and extinction is a central question in evolutionary biology. Birth–death models provide a mathematical framework for estimating these rates from phylogenetic trees and are widely used to study the tempo of diversification. Reliable estimates of speciation (*λ*) and extinction (*µ*) rates form the basis for downstream analyses, such as testing for shifts in diversification across lineages or through time (e.g., Nee et al. 1992; Rabosky 2006, 2009; Bokma 2008; Rabosky and Lovette 2008; Morlon et al. 2011; Etienne et al. 2012), identifying lineage-specific heterogeneity (e.g., Alfaro et al. 2009; Rabosky 2014; Maliet et al. 2014; Vasconcelos et al. 2022a), and linking diversification patterns to ecological and/or trait data (e.g., Slowinski and Guyer 1993; Maddison et al. 2007; Beaulieu and O’Meara 2016).

While there are various ways to estimate rates from these models, the most common is maximum likelihood (or related Bayesian approaches). Maximum likelihood has many advantages, such as consistency with increasing sample size and efficient use of information in the data. However, likelihood estimators are often biased at small sample sizes. For many questions, sample sizes are necessarily limited: there are only so many species of living whales, so any approach for understanding modern whale diversificaton has a small number of data points. Moreover, many popular approaches (e.g., Medusa, Alfaro et al. 2009; BAMM, Rabosky 2014; ClaDS, Maliet et al. 2014; MiSSE, Vasconcelos et al. 2022a) subdivide a tree into smaller parts, each with its own estimates of rates.

We have become concerned with several potential sources of bias that can influence inference. One concern is *statistical bias* – that is, systematic deviations in the expectations of birth and death rate estimators from their true generating values. Such biases produce apparent trends in diversification that are artifacts of the estimation procedure rather than features of the underlying process, especially with small sample sizes. For instance, the commonly used Yule birth rate estimator (Yule 1924; Nee 2001) systematically underestimates speciation rates (O’Meara and Beaulieu 2024). A second concern is *structural bias*, arising from how likelihoods are conditioned and how trees of particular sizes are treated. Many formulations assume survival of the crown clade (e.g., Stadler 2013), but in practice trees of particular sizes are often excluded from analysis. Specifically, two-taxon trees (*n* = 2) are often excluded, either because the likelihood is not defined or because such clades are rarely, if ever, subject to rate estimation. This implicit filtering introduces an additional layer of conditioning that could alter the distribution of observed clades and, in turn, affect parameter estimates. A third potential source is *ascertainment bias*, which is when researchers preferentially select unusual or noteworthy clades. While this does not bias estimates within those clades, it can limit how representative they are of broader diversification patterns. Finally, ignoring uncertainty in measurements introduces its own biases; for more on that, see O’Meara and Beaulieu (2024).

In this paper, we examine the mathematical underpinnings of the first two potential biases and propose solutions. We begin by analyzing the behavior of rate estimators on the smallest possible clades, demonstrating why *n* = 2 trees lack sufficient information to identify both speciation and extinction separately. Throughout, we refer to complete two-taxon trees as cherry trees – that is, rooted trees with only two terminal edges. This usage distinguishes them from internal “cherries” (pairs of sister taxa within larger trees; see McKenzie and Steel 2000) and emphasizes their role as minimal cases in diversification models. We then derive the appropriate conditioning for likelihoods when cherry trees are excluded from empirical datasets. Building on this, we then revisit the Yule estimator and derive an analytical bias correction, clarifying how an earlier empirical correction from our previous work relates to theory.

Finally, we extend these insights to the general birth–death model, where extinction complicates analytical results, and use symbolic regression to obtain practical bias corrections. Together, these results highlight the importance of careful statistical treatment when estimating diversification rates and provide a framework for reducing biases in diversification rate analyses when dealing with small trees or subtrees.

## 2 Likelihood functions and the problem of cherry trees

With the Yule model of phylogenetic tree growth, there is only a birth rate, *λ*, and no extinction (*µ*=0). This means that once a lineage arises, it persists indefinitely, without the possibility of going extinct. Consequently, every branch of the tree represents a successful speciation event without subsequent loss. The Yule model is often considered a framework that captures unbounded species diversification dynamics. The maximum likelihood estimate of *λ* can be calculated directly using the formula derived in Nee (2001):

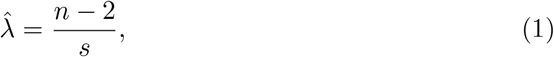

where *n* is the number of tips in the tree and *s* represents the sum of all *x*_*i*_ edge lengths in the tree [i.e., Σ(*x*_*i*_)]. Notice that when *n* = 2 estimates of 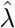 become zero, which is intuitive. In the absence of extinction, initializing a tree at *n* = 2 that remains unchanged after some time interval implies that no additional speciation events occurred, which is a speciation rate of zero.

With a standard birth-death model, where we assume that extinction is *µ* > 0, when estimating rates on a cherry tree, we have more parameters than observed speciation events: there is simply not enough information in them to uniquely estimate both *λ* and *µ* (see Appendix A). However, there are more immediate and practical considerations. As discussed by Stadler (2013), most implementations of the constant birth-death model rely on the same underlying likelihood function. Specifically, many use a form of,

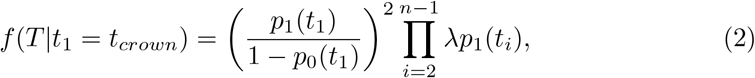

which assumes that the process is conditioned on the survival of the crown – that is, that at least one lineage persists from each of the daughter lineages of the root (see Stadler 2013). Without delving into the specifics of how to calculate *p*_1_ and *p*_0_ in the above expression, we direct attention to the product term on the right-hand side. This term iterates over a vector of *n* − 1 speciation times from a phylogeny with *n* species, but the indexing begins at *i* = 2. In other words, the product excludes the crown node, which is handled separately by the term on the left. When the tree is a cherry tree (*n* = 2), the vector of speciation times is empty and the term of the product is undefined. In practice, this means that the likelihood cannot be computed for two-taxon trees using this formulation (or any variant thereof; see Stadler 2013), and therefore implicitly exclude such trees by assuming that tree sets contain only *n* ≥ 3 trees.

The exclusion of two-taxon trees has important consequences, especially with young clades. To illustrate, we simulated 100,000 trees under a constant-rate birthdeath process (*λ*=0.1, *µ*=0.05), estimating both *λ* and *µ* using the likelihood function of Stadler (2013) across a range of clade ages. As expected, estimates based on all surviving trees (but excluding those with only two tips) show a clear trend: younger clades exhibit upwardly biased diversification rates (Table 1).

**Table 1:**
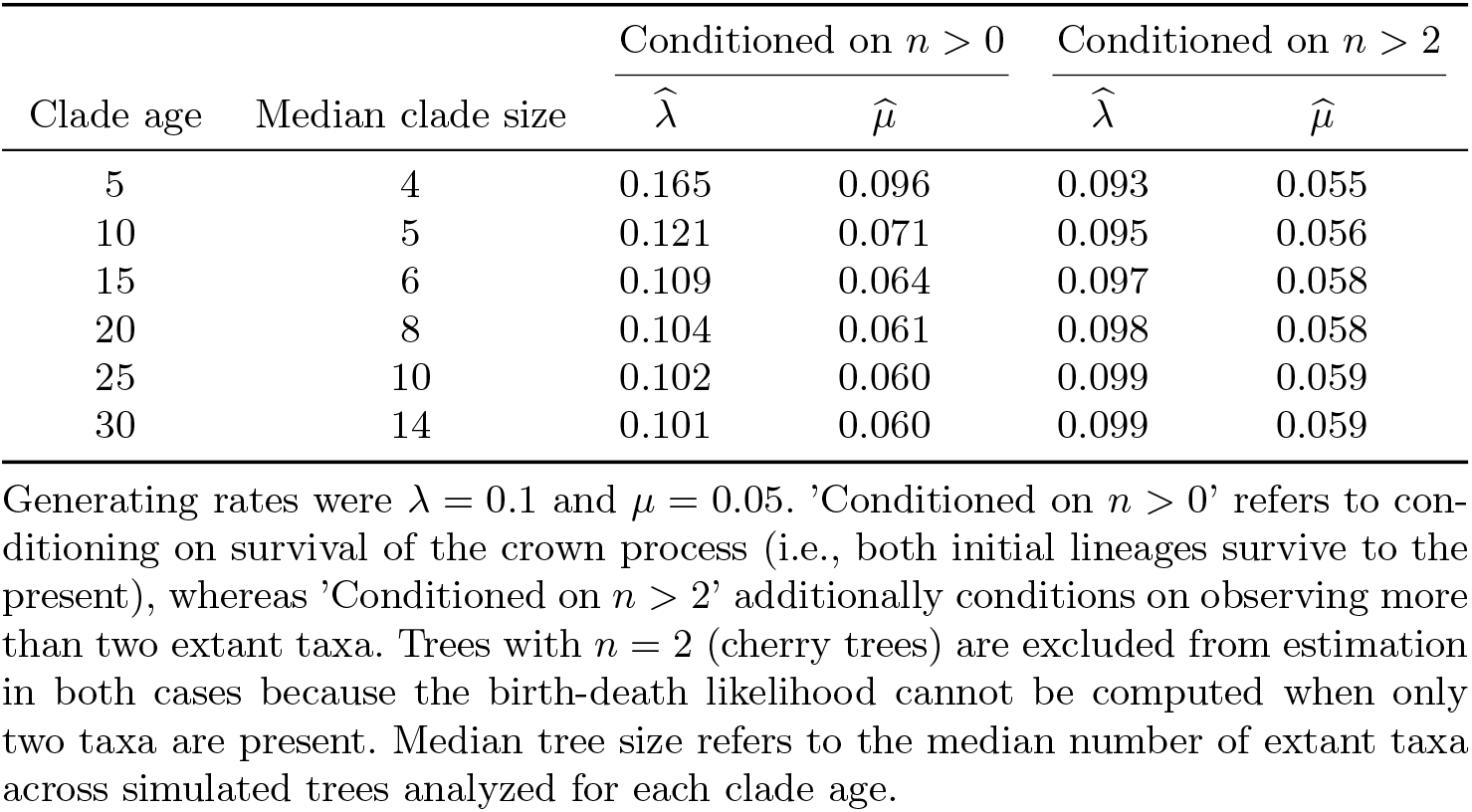
Summary of estimates of speciation and extinction rates using the likelihood formula of Stadler (2013).

| Clade age | Median clade size | Conditioned on $n > 0$ | | Conditioned on $n > 2$ | |
| --- | --- | --- | --- | --- | --- |
| | | $\hat{\lambda}$ | $\hat{\mu}$ | $\hat{\lambda}$ | $\hat{\mu}$ |
| 5 | 4 | 0.165 | 0.096 | 0.093 | 0.055 |
| 10 | 5 | 0.121 | 0.071 | 0.095 | 0.056 |
| 15 | 6 | 0.109 | 0.064 | 0.097 | 0.058 |
| 20 | 8 | 0.104 | 0.061 | 0.098 | 0.058 |
| 25 | 10 | 0.102 | 0.060 | 0.099 | 0.059 |
| 30 | 14 | 0.101 | 0.060 | 0.099 | 0.059 |

Generating rates were $\lambda = 0.1$ and $\mu = 0.05$ . 'Conditioned on $n > 0$ ' refers to conditioning on survival of the crown process (i.e., both initial lineages survive to the present), whereas 'Conditioned on $n > 2$ ' additionally conditions on observing more than two extant taxa. Trees with $n = 2$ (cherry trees) are excluded from estimation in both cases because the birth-death likelihood cannot be computed when only two taxa are present. Median tree size refers to the median number of extant taxa across simulated trees analyzed for each clade age.

We note that this issue does not arise when using alternative likelihood frameworks that handle small trees more directly. For example, the approach developed by Maddison et al. (2007) that is implemented in state-dependent speciation and extinction models (SSE) does not rely on the product over internal nodes and thus can accommodate trees with as few as two taxa. The unconditioned likelihood under this formulation is,

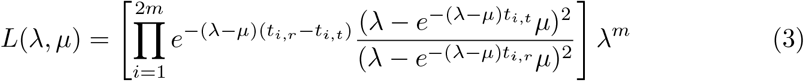

where *m* is the number of internal nodes, and thus 2*m* is the number of branches in the tree, *T* is the total height of the tree, and finally *t*_*i,t*_ and *t*_*i,r*_ reflect the tipward and rootward ages of node *i*. Because the likelihood is expressed as a product over branches rather than over speciation events, the indexing begins at *i* = 1 even for the smallest possible tree. As a result, trees with *n* = 2 taxa (*m* = 1) are naturally accommodated under this formulation.

However, this raises an important question: even if a likelihood can be evaluated for *n* = 2, does such a tree contain enough information to estimate both speciation and extinction rates? Analytical results show that it does not – cherry trees simply lack the complexity needed to disentangle these two processes, even when the likelihood is defined (see Appendix A). We therefore recommend excluding two-taxon trees (or subtrees) from analyses, but conditioning the likelihood accordingly to account for their absence.

## 3 Conditioned likelihood for *n* > 2 censored tree sets

Two-taxon trees contain little information for estimating speciation and extinction rates, and in some likelihood formulations their probability cannot be computed directly. In practice, such trees are often excluded – sometimes explicitly by imposing a minimum clade size, and sometimes implicitly because likelihood terms are undefined. This exclusion introduces an additional form of conditioning beyond the usual requirement that the crown lineage survives to the present (i.e., Stadler 2013). In other words, inference is restricted to clades with more than two extant taxa. This second layer of conditioning is nontrivial because the probability of observing a two-taxon tree depends on both the underlying model and the clade’s age, and therefore varies under varying birth-death dynamics. To account for this, we derive the appropriate conditioning term when datasets are censored to exclude *n* ≤ 2 trees, under a general birth–death process and its two special cases: the Yule process (*µ* = 0) and the critical branching process (*λ* = *µ*).

### 3.1 Birth-death model (*λ ≠ µ*)

We begin with Eq. 3 which is the unconditioned likelihood equation of Maddison et al. (2007). Conditioning on the existence of the crown node with two surviving lineages requires scaling the likelihood by the probability of survival. The probability that a single lineage goes extinct after time, *t* is:

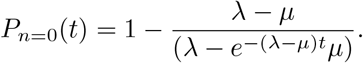

Thus, the probability of survival of the two descendants of the root is, [1 − *P*_*n*=0_(*t*)]^2^. Accounting for the initial speciation event at the crown reduces the exponent of *λ* from *m* to *m* − 1. Together, the likelihood conditioned on crown survival becomes,

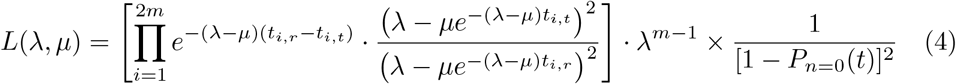

We now further condition on observing trees with more than two extant taxa. Following Stadler (2013), the probability that a single lineage produces exactly one descendant after time, *t*, is

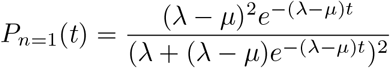

For a crown with two descendants, the probability that both lineages survive but each produces exactly one extant descendant tip (i.e., the tree has exactly two tips) is *P*_*n*=1_(*t*)^2^. Therefore, the probability that the crown survives and produces more than two extant taxa is obtained by subtracting this event from the probability of crown survival,

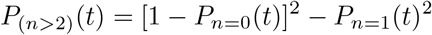

Conditioning the likelihood on observing more than two extant taxa then yields:

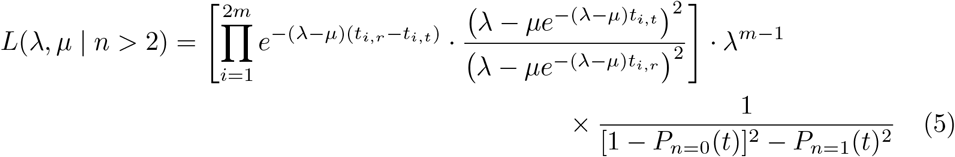

Having derived the appropriate conditioning for datasets where *n* = 2 trees are excluded, we next evaluated whether incorporating this adjustment corrects the systematic bias observed in simulations (Table 1). When we return to the simulated datasets, the effect of applying this additional conditioning is clear. If we modify the likelihood so that it reflects not only the probability that the crown lineage at time *t* survives to the present, but also that it gives rise to a clade with *n* > 2, the upward trend in rate estimates disappears. In fact, the estimates for *λ* become slightly downward biased compared to the true generating parameters (Table 1).

### 3.2 Special case 1: Yule model (*µ* = 0)

Following Nee (2001, Eq. 3), the likelihood of an observed tree under a Yule model is defined as,

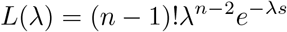

where, again, the term *s* represents the sum of all *x*_*i*_ edge lengths in the tree. To condition the log-likelihood on the observed tree by the probability of producing any tree with more than two taxa. In other words, *P* (*n* > 2) represents the probability that a tree of the exact same age and speciation rate would have more than two taxa. To condition on *n* > 2, we start with the probability of *n* lineages, given *λ* and *t*, from Nee (2001),

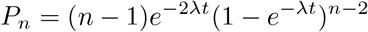

where *t* represents the length from root to tip. Note that the use of *t* is to distinguish it from *s* in the log-likelihood. If we assume that *n* = 2, this equation simplifies to:

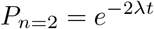

Therefore, the probability that a tree has *n* > 2, given *λ* and *t* is

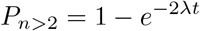

and the conditioned likelihood is calculated as:

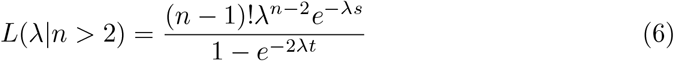

### 3.3 Special case 2: Critical branching process (*λ* = *µ*)

With the general birth-death model, the probability of a phylogeny depends on both *λ* and *µ*. However, when *λ* = *µ* the contribution of *µ* to the likelihood becomes indistinguishable from that of *λ*. Specifically, both speciation and extinction events contribute equally to the branching patterns in the tree, and their exact roles in determining the timing and structure of the tree are indistinguishable from each other. As a result, the likelihood function is simplified and depends only on a single parameter, *λ*. Following the derivation provided by Maddison et al. (2007), the unconditioned likelihood of an observed tree under the critical branching process model:

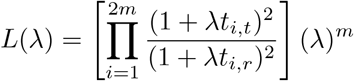

Bailey (1964, Eq. 8.53) derived the probability of observing *n* species at time, *t*, under a critical branching process (i.e., when *λ* = *µ*):

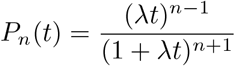

As with the general birth-death model, we are interested in the probability that a crown group of age, *t*, both survives (i.e., both initial lineages leave descendants) and produces more than two tips. The probability of extinction of a single lineage after time, *t*, is calculated as (Maddison et al. 2007):

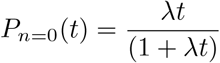

Thus, the probability of survival of the two descendants of the root is,

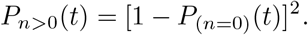

The probability of generating exactly two tips (i.e., both lineages yield exactly one descendant) is:

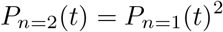

Therefore, the joint probability of crown survival and tree size *n* > 2 is:

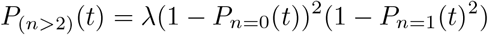

Accounting for the speciation event at the crown, the new conditioned likelihood then becomes:

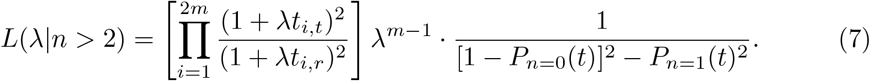

## 4 Correcting estimator bias

In a recent publication (O’Meara and Beaulieu 2024), we showed that the Yule estimator is actually biased *downwards* as a function of time – that is, as time approaches the present, or across short time intervals, rate estimates increasingly fall below the generating value. We suspect that this phenomenon has gone largely unnoticed because trees that fail to speciate (trees of *n* = 2) after time *t* are almost always discarded. Excluding these trees make it seem like rates increase dramatically towards the present or across shorter time intervals. In our study, we used a bias “correction” of *n/*(*n*−1) that produced a remarkably constant trend through time when the zero rates are included. What was not clear from the paper is that this correction was actually derived empir-ically from simulation results (we simply showed that the naive estimator was indeed biased). As we detail below, the proper correction term is (*n* − 1)*/*(*n* − 2).

By contrast, analytically deriving bias corrections for birth-death models is con-siderably more difficult, even though both *λ* and *µ* appear downwardly biased across short timescales (Table 1). The presence of extinction introduces nonlinear dependencies between parameters, making it intractable to isolate closed-form expectations of the estimators. This complexity arises because the likelihood under a birth-death process depends jointly on speciation (*λ*) and extinction (*µ*) and becomes even more complex when it is conditioned on survival and a minimum number of extant taxa. As a result, while exact analytical corrections are not readily available, we used symbolic regression to search for *approximate* transformations of the estimated parameters that systematically reduce bias. To validate this approach, we first applied it to trees simulated under a Yule process and confirmed that symbolic regression recovered the known analytical correction. We then extend this method to estimates from trees simulated under a wide range of birth-death processes.

### 4.1 Bias correction for the Yule model

A key insight from Steel and Mooers (2010) is that, under a Yule process conditioned on the observed number of tips, the expected average edge length is 1*/*(2*λ*), as opposed to 1*/λ*, because each speciation creates two lineages. We begin by restating the standard Yule likelihood formula (Nee 2001) from Eq. 1, where the MLE estimator of *λ* is computed as

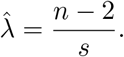

Let *s* = Σ*x*_*i*_ denote the total branch length in a bifurcating tree with 2*n*−2 edges and use the edge-length expectation *E*[*x*_*i*_] = 1*/*(2*λ*), then 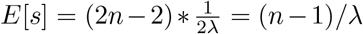. Here, the expectation is taken conditional on the observed number of tips *n*, so that the numerator *n*−2 is fixed and all stochasticity lies in the total tree length *s*. Plugging this into the expectation for 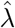 yields

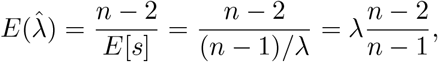

so 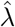 is downward biased. Because *n* is fixed under the likelihood, this expression gives the conditional expectation of the Yule maximum-likelihood estimator and does not assume a general equivalence between *E*[*X/Y* ] and *E*[*X*]*/E*[*Y* ]. To derive the bias of the MLE estimate of *λ*, we then subtracted *λ* from 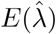:

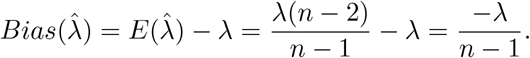

To construct an unbiased estimator from this, we found a multiplier *c* that satisfies

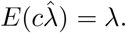

Substituting 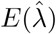,

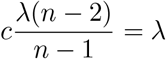

and then solving for *c* we obtain the unbiased estimator as,

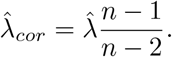

Empirically, O’Meara and Beaulieu (2024) showed that applying a *n/*(*n*−1) correction to the Yule estimator worked well in removing the bias. This apparent discrepancy can be understood by considering an alternative conditioning. If instead of conditioning on the number of tips we consider a Yule process that grows for a fixed time, *t*, and the tip number *n* varies, Kendall (1949) derived the variance of the estimator as

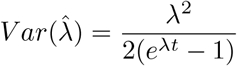

This expression describes variation among replicate trees of equal age *t* but differing sizes, rather than among trees of fixed *n*. Since the bias-corrected estimator is obtained by multiplying the original by a constant factor, its variance scales by the square of that factor. Thus,

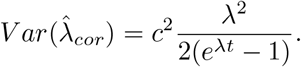

Although this variance pertains to the fixed-time case, it illustrates how different small-sample corrections 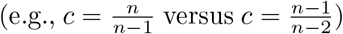 affect the bias–variance trade-off: the former yields slightly lower variance but introduces bias under the fixed *n* likelihood.

### 4.2 Bias correction for general birth–death models

While analytical bias corrections are tractable for the Yule process, they are not for the general birth–death process where extinction is present. To address this, we developed a symbolic regression approach to search for approximate transformations of the estimated parameters that systematically reduce bias. Before applying this method to general birth–death models, we first validated it under the Yule process to ensure that it could recover known analytical corrections (see above). We then extended the approach to simulations under a broad range of birth–death scenarios, where closed-form expectations are unavailable.

#### 4.2.1 Validating symbolic regression with the Yule model

To validate the symbolic regression approach, we simulated 250,000 Yule trees that span a wide range of clade ages and speciation rates. To do so, we first sampled 250,000 pairs of points for clade age and speciation rate (*λ*) using a Latin hypercube sampling design, drawing each variable independently from a uniform distribution over the interval [0,5] and [0,0.5], respectively. For each combination of clade age and speciation rate, trees were simulated and those with *n* = 2 were discarded. This yielded 140,449 trees, for which rates were estimated using the standard Yule likelihood conditioned on *n* > 2.

Using the R package `gramEvol` (Noorian et al. 2016) we defined a grammar that allows transformations and combinations of relevant variables using basic algebraic operations and mathematical functions. We included the following variables: 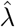 is the likelihood estimate for speciation conditioning only on crown survival; *n* is the number of taxa in a given tree; (*n* − 1)*/*(*n* − 2) and *n/*(*n* − 1) are potential sample size bias correction functions; and a placeholder for constants (see Appendix B). To ensure dimensional consistency and interpretability, we retained only expressions in which the correction took the multiplicative form 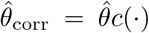, where *c* is a dimension-less function of sample size and dimensionless combinations of estimated rates and constants. This restriction prevents unit-inconsistent expressions while maintaining sufficient flexibility to correct bias. Each symbolic expression generated by the grammar was evaluated by comparing its predicted output across simulated trees to the known generating speciation rates. Specifically, the true rates were treated as predictors and the output of the symbolic expressions as responses in a robust linear regression. We settled on using a robust linear model with the Huber loss function to downweight the influence of outliers when evaluating how closely each expression recovered the true generating values: various other ways of creating loss functions, like using RMSE, were very sensitive to some of the extremely bad estimates from small trees. Transformations were selected based on minimizing squared deviations from the 1:1 relationship between predicted and true rates, identifying those that systematically reduced bias in the estimated parameters.

To better explore the expression space, we implemented a custom exhaustive search function instead of using the default evolutionary search within `gramEvol`, as preliminary runs indicated that the evolutionary algorithm did not reliably recover the lowest-scoring expressions under our grammar and depth constraints. This exhaustive search enumerated all admissible expressions up to a user-defined maximum depth (`max.depth` = 4); deeper expressions were not considered because the size of the resulting expression space grows combinatorially and becomes computationally intractable. To penalize for model complexity, we added the term Ω = *α* · depth to each cost score, where depth is the depth of the expression tree and *α* is a regularization parameter controlling the strength of the penalty. We tested a range of *α* values from 0 to 0.02 to determine how strongly one must penalize complexity before a simpler expression overtakes the true model; notably, *α*=0.02 corresponds approximately to the loss score of the uncorrected 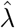, providing a natural upper bound for the regular-ization strength. Smaller values of *α* indicate that only mild penalization is needed to favor simpler expressions, whereas larger values imply that even under strong penalization the more complex model remains the better fit. This approach allowed us to assess the robustness of the analytically derived correction under increasing levels of regularization.

Table 2 summarizes the raw scores across the top five expressions returned. For most of the *α* range (*α* ∈ [0, 0.0169], covering 84.6% of the grid), the analytically derived correction 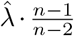 achieved the lowest penalized cost score, with a raw fit error nearly two orders of magnitude smaller than that of the uncorrected 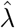. Only when *α* exceeded 0.0169, where the complexity penalty was nearly as large as the fit score itself, did the uncorrected 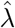 briefly become favored (15.4% of the grid). However, the advantage was driven purely by penalization, not fit. Thus, the analytically derived correction remained the best-supported expression across a wide range of penalty weights, demonstrating that it is robust to regularization and not simply favored under weak complexity penalties.

**Table 2:** Ranking of symbolic expressions for corrections to the Yule estimator.

| Rank | Expression | Depth | Raw Score |
| --- | --- | --- | --- |
| 1 | $\hat{\lambda} \cdot \frac{n-1}{n-2}$ | 2 | 0.00013 |
| 2 | $\hat{\lambda} \cdot \frac{n}{n-1}$ | 2 | 0.00066 |
| 3 | $\hat{\lambda} \cdot \frac{(n-1)^2}{n(n-2)}$ | 3 | 0.01202 |
| 4 | $\hat{\lambda} \cdot \frac{n}{n-2}$ | 3 | 0.01465 |
| 5 | $\hat{\lambda}$ | 3 | 0.01704 |

Finally, as shown in Figure 1, the uncorrected estimates systematically underestimated the true value, with the fitted slopes falling below one. In contrast, the corrected estimates aligned with the 1:1 line throughout the entire range of simulated values, confirming that the analytically derived correction of the naive estimator not only eliminates bias but also produces robust and accurate speciation rate estimates.

**Fig. 1:**
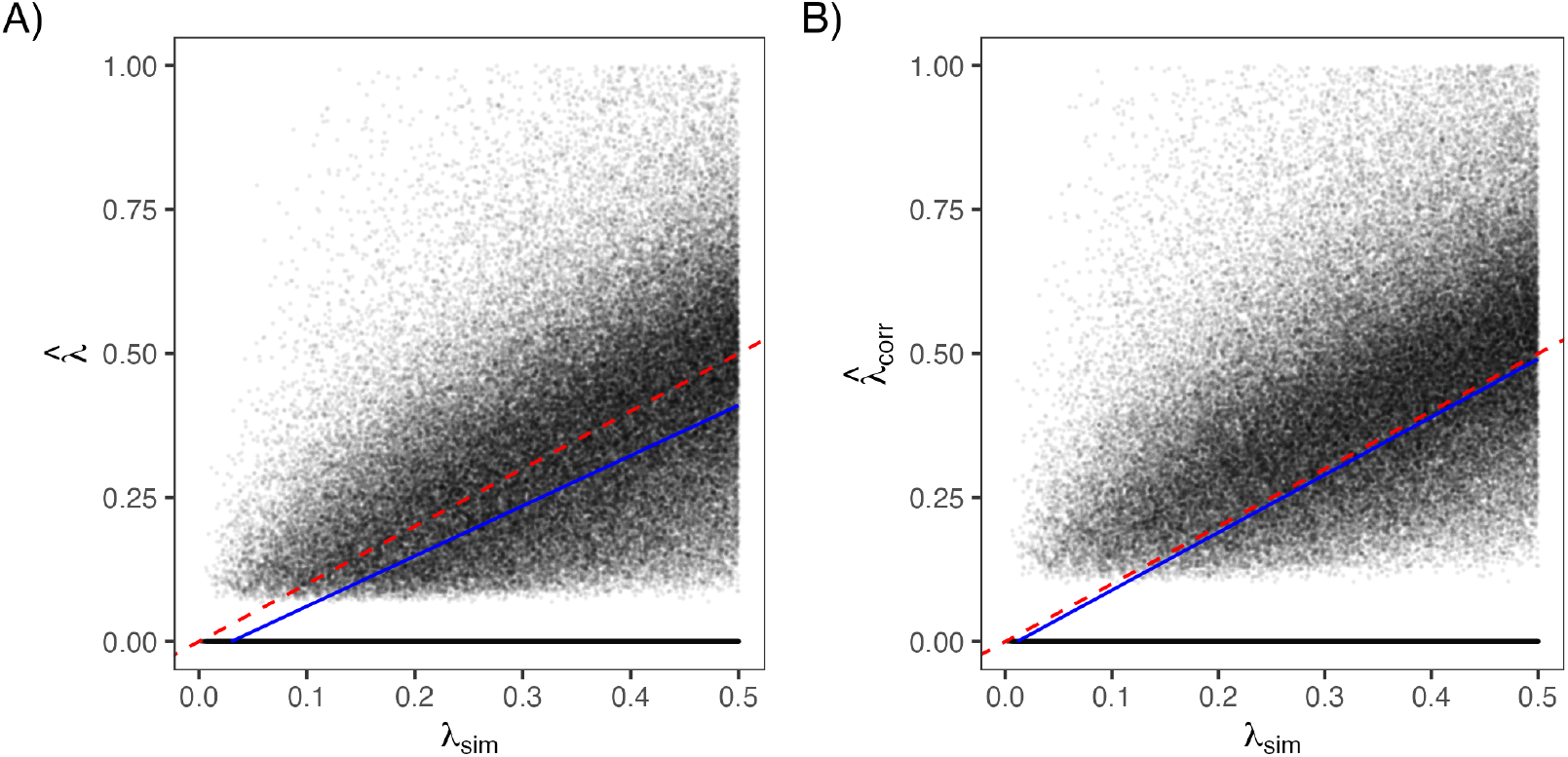
Effect of bias correction on estimates of the speciation rate *λ* in Yule-simulated trees. Each panel shows estimated values of *λ* plotted against the true simulated value *λ*_sim_ across nearly 150,000 trees. (A) Uncorrected estimates 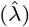 systematically underestimate the true value. (B) Applying the correction factor 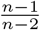 to 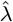 improves accuracy, bringing estimates closer to the 1:1 line (dashed red). Blue lines show robust linear model fits using Huber M-estimation.

#### 4.2.2 Extending to general birth–death processes

Having confirmed that symbolic regression recovers the known Yule correction, we next applied the method to trees simulated under a wide range of birth–death processes. We sampled 500,000 sets of points for clade age, speciation rate (*λ*), and extinction fraction (*ε* = *µ/λ*), again, using a Latin hypercube sampling design, drawing each variable independently from a uniform distribution over the interval [0,10], [0,1], and [0,1], respectively. Note that the simulated value for *µ* was based on multiplying *λ* by *ε*. Trees were then simulated for each set of clade age, speciation rate, and extinction rate, discarding any tree where *n* ≤ 2. Speciation (*λ*) and extinction (*µ*) rates were then estimated using the standard birth–death likelihood conditioned on *n* > 2. However, rather than estimating *λ* and *µ* directly, we first estimated two composite parameters – turnover (*τ* = *λ* + *µ*) and extinction fraction (*ε* = *µ/λ*) – and then back-transformed these values to recover estimates of *λ* and *µ*. To ensure that maximum likelihood estimates reflected genuine maxima, we excluded simulation replicates in which the variance–covariance matrix derived from the Hessian returned negative variance for either parameter. In total, this resulted in 346,957 trees from which we were able to obtain robust estimates of *λ* and *µ*.

The symbolic regression was then used to identify transformations that removed the downward biases exhibited by *λ* and *µ* with respect to their known generating values (see Appendix B). However, unlike with the Yule simulations, we did not have an analytically known “best” expression for the birth–death case. Instead, we identified the leading expressions by systematically varying the regularization weight *α* across a dense grid of 5,000 points. For each *α*, we computed the total penalized score and recorded the expression with the lowest value. Whenever the highest-ranked expression changed, we defined an interval of *α* values over which the previous expression remained optimal, along with summary metrics including the minimum and maximum *α* and Ω at the endpoints. This procedure allowed us to trace how expression rankings shifted as the penalty for complexity increased, distinguishing corrections that were robust across broad *α* intervals from those that appeared only under extreme regularization. The search range for *λ* was *α* ∈ [0, 10^−3^] and *α* ∈ [0, 10^−1^] for *µ*. These bounds ensured that all regions with non-negligible penalized score were explored while avoiding oversampling in irrelevant ranges.

Table 3 summarizes the main results of symbolic regression for *λ* and *µ* under the birth–death model. For *λ*, the expression 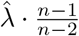 consistently achieved the lowest penalized cost over nearly the entire range of regularization weights. This correction reduced the raw fitting error by more than an order of magnitude relative to the uncorrected 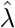, and remained favored until the complexity penalty became comparable in magnitude to the fit score itself (*α* ≈ 0.0015). Beyond this point, the uncorrected 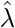 was marginally preferred, though the improvement was driven solely by penalization rather than by fit. Thus, a correction identical to the one found for *λ* under the Yule process also emerged as the most stable and parsimonious adjustment in the birth–death case.

**Table 3:** Candidate expressions for speciation (*λ*) and extinction (*µ*) rates under a birth–death model, ranked by their raw score.

| Parameter | Expression | Depth | Raw Score | $\alpha_{\min}$ | $\alpha_{\max}$ | $\Omega_{\min}$ | $\Omega_{\max}$ |
| --- | --- | --- | --- | --- | --- | --- | --- |
| $\lambda$ | $\hat{\lambda} \cdot \frac{n-1}{n-2}$ | 2 | 0.00041 | 0.00000 | 0.00148 | 0.00000 | 0.00297 |
| $\lambda$ | $\hat{\lambda}$ | 1 | 0.00189 | 0.00148 | 0.00500 | 0.00148 | 0.00500 |
| $\mu$ | $\hat{\mu} \cdot (\frac{n}{n-1} + \hat{\varepsilon})$ | 4 | 0.00665 | 0.00000 | 0.01164 | 0.00000 | 0.04625 |
| $\mu$ | $\hat{\mu} \cdot 2 \frac{n-2}{n-1}$ | 3 | 0.01827 | 0.01164 | 0.02667 | 0.03493 | 0.07999 |
| $\mu$ | $\hat{\mu} * 2$ | 2 | 0.04493 | 0.02667 | 0.10000 | 0.05333 | 0.20000 |

For *µ*, the symbolic regression identified a small set of multiplicative corrections whose structure varied systematically with the regularization penalty. Across low penalty values (*α* ∈ [0, 0.0116]) was 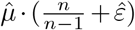, indicating that the downward bias in *µ* bias depends jointly on sample size and the estimated extinction fraction. Unlike the speciation rate, where bias is driven primarily by sample size effects, this result reflects the intrinsic coupling between *λ* and *µ* in the birth–death likelihood: uncertainty in 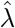 propagates directly into estimates of 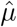 through 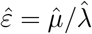. As the regularization penalty increased, simpler corrections were favored, first 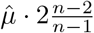 and then 2 · 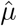. These latter expressions capture the dominant direction of bias but ignore its dependence on *ε*. Taken together, these results show that while bias in *µ* can be partially mitigated by simply multiplication of constant, the most accurate corrections explicitly incorporate both sample size effects and extinction fraction, underscoring, perhaps unsurprisingly, that *µ* bias is structurally more complex than *λ* bias under birth-death models.

#### 4.2.3 Bias corrections and derived diversification parameters

Having derived bias corrections for *λ* and *µ*, we next examined how these adjustments affect other diversification parameters – specifically, turnover (*λ* + *µ*) and net diversification (*λ* − *µ*) – that are commonly used to summarize macroevolutionary dynamics. To evaluate whether the symbolic regression-based corrections for *λ* and *µ* fully removed systematic bias, we also derived expectation-based corrections by regressing the corrected rate estimates on their generating values across simulations. These expectation-based corrections serve as an empirical calibration step, whereas symbolic regression targets bias in the estimators, the regression-based mapping removes the small, approximately linear residual bias that remains when corrected estimates are compared directly to their true values in simulation. This produces linear mappings of the form

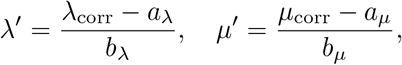

where *a* and *b* are the intercept and slope from the corresponding regression fits. These transformations recenter the corrected estimates on their expected values under the simulation design and are used here to assess how residual bias propagates into composite diversification parameters. Turnover and net diversification were then recomputed using both the directly corrected and expectation-based rates.

When turnover was calculated directly from the corrected *λ* and *µ*, the relationship with simulated values was nearly unbiased, with an intercept near zero (-0.003) and a slope slightly above one (1.086). This reflects the partial cancellation of small residual biases in *λ* and *µ*, where both corrected estimators slightly under predict at higher values but in similar proportion. Using the expectation-based corrections yielded a comparable fit (slope = 1.049, intercept = -0.037), indicating that turnover is largely robust to modest differences in the individual rate corrections (Fig. 2).

**Fig. 2:**
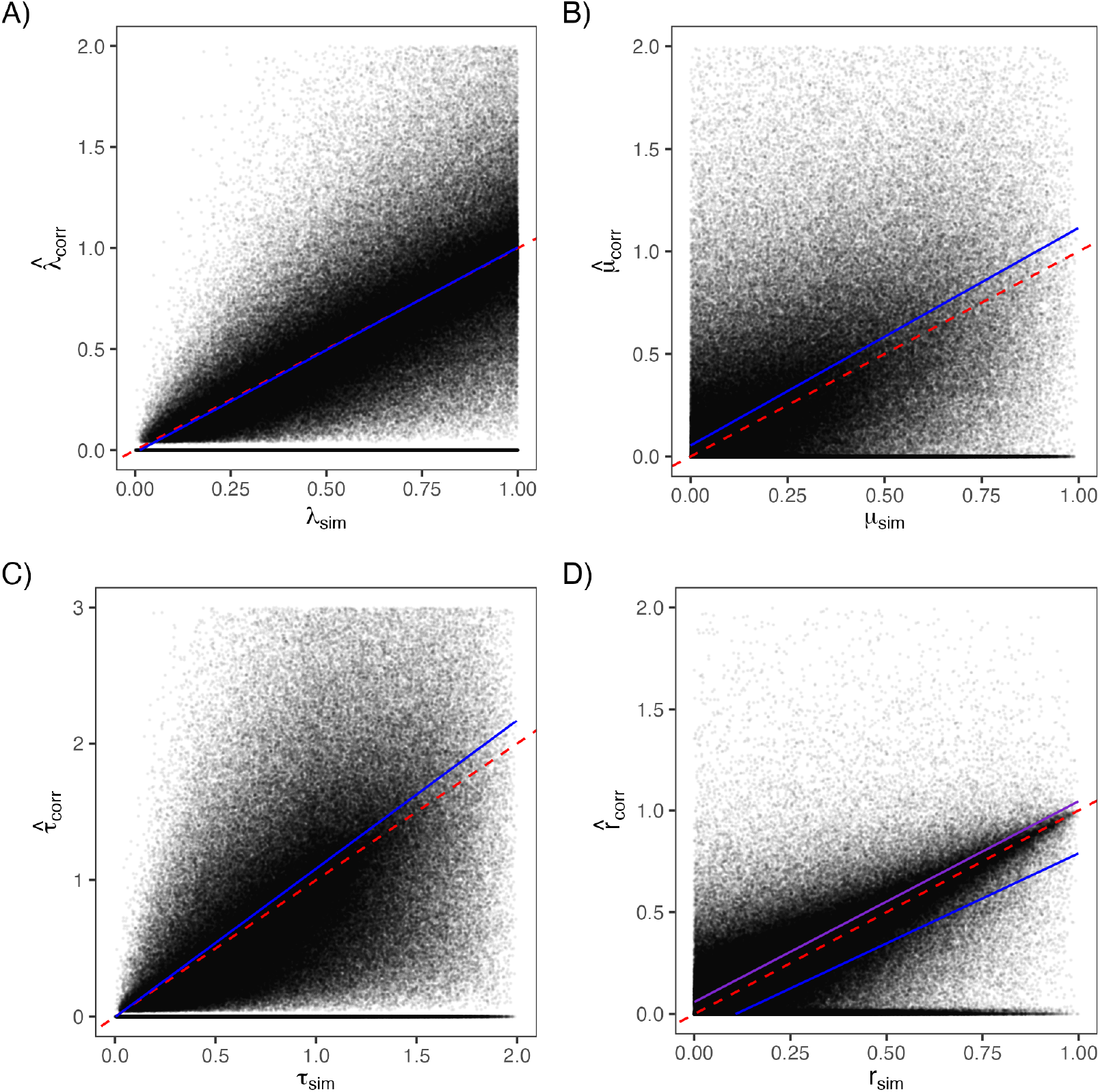
Relationships between simulated (“sim”) and corrected estimates of (A) speciation rate, (B) extinction rate, (C) turnover rate, and (D) net diversification. Each panel shows the empirical regression line (dark blue) and the 1:1 line (dashed red). The close alignment between the two lines in panels (A-C) indicates that bias corrections for *λ* and *µ* largely remove systematic deviation, while panel (D) shows that net diversification remains slightly underestimated at high true values due to small asymmetries between *λ* and *µ* corrections; however, when we correct net diversification directly regression line (purple) is much closer to the 1:1 line.

In contrast, net diversification (*λ* − *µ*) remained more sensitive to differential bias between *λ* and *µ*. When computed directly from the corrected rates, the slope was notably below one (0.884), and the intercept negative (-0.095), reflecting persistent underestimation at higher true net-diversification values. Applying the expectationbased correction slightly improved this relationship (slope = 0.874, intercept = -0.019) but did not fully remove the bias (Fig. 2).

Symbolic regression provided insight into this behavior. Unlike speciation, whose bias is well captured by sample size, both extinction and net diversification were best corrected by expressions that explicitly depended on the estimated extinction fraction. The optimal correction for net diversification matched that for *µ*, indicating that the systematic error in 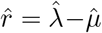 is dominated by the same coupling between extinction and speciation that drives bias in 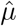. In effect, although both 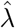 and 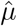 are estimated without prior correction, systematic error in 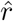 is dominated by the same extinction-fractiondependent bias that affects 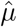, leading the optimal net-diversification correction to match that for *µ*. Applying the symbolic-regression–derived correction substantially improved agreement with the true net diversification values, producing a nearly one-to-one relationship (slope = 0.986), although with a small positive intercept (0.059), indicating a slight tendency to overestimate net diversification across the entire range of generating values (Fig. 2).

On the whole, these results show that turnover behaves as a stable composite of *λ* and *µ*, whereas net diversification is more strongly affected by small asymmetries in their respective biases, resulting in a consistent flattening of its regression slope. This is primarily due to *µ* being slightly overestimated relative to *λ*. These patterns are consistent with the algebraic expectations derived from the corrected estimators, where net diversification inherits asymmetric bias from the difference between 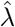 and 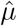 expectations. Because the magnitude of this difference increases with true net diversification, the slope of estimated versus simulated values remains systematically below one, even when both underlying rate corrections are nearly unbiased.

## 5 Discussion

The results presented here highlight additional statistical and structural sources of bias in the estimation of diversification rates under the birth-death framework. With respect to estimator bias, we showed analytically that for the Yule process (*µ* = 0) the maximum likelihood estimator for *λ* is biased downward. Our empirically driven approach to approximate the downward bias for *λ* and *µ* (when *µ* > 0) in the birthdeath model identified functional expressions that minimized the error between the generating and estimated values. The best-performing correction for *λ* in a birth-death model was identical to the correction in the Yule model. Although extinction alters the distribution of observed tree sizes by increasing the probability of small clades, it does not change the sample size bias of the speciation rate estimator once the likelihood is properly conditioned on survival and on observing more than two extant taxa. In contrast, *µ* exhibited a more complex pattern, where the bias depended jointly on both sample size and extinction fraction (*µ/λ*). The recommended corrections for *λ, µ*, turnover, and net diversification are summarized in Table 4.

**Table 4:** Summary of recommended bias corrections under the birth–death model.

| Parameter | Recommended correction |
| --- | --- |
| Speciation ($\lambda$) | $\hat{\lambda}_{\text{corr}} = \hat{\lambda} \cdot \frac{n-1}{n-2}$ |
| Extinction ($\mu$) | $\hat{\mu}_{\text{corr}} = \hat{\mu} \cdot \left( \frac{n}{n-1} + \hat{\varepsilon} \right)$ |
| Turnover ($\tau = \lambda + \mu$) | $\hat{\tau}_{\text{corr}} = \hat{\lambda}_{\text{corr}} + \hat{\mu}_{\text{corr}}$ |
| Net diversification ($r = \lambda - \mu$) | $\hat{r}_{\text{corr}} = \hat{r} \cdot \left( \frac{n}{n-1} + \hat{\varepsilon} \right)$ |

Both *λ* and *µ* are underestimated by their maximum likelihood estimators, but the bias is not symmetrical. After applying the correction, *λ* closely follows the 1:1 relationship with its generating values, whereas *µ* tends to remain slightly overestimated. Because turnover is the sum of *λ* and *µ* it seems that the opposing biases in the two parameters largely offset each other, producing nearly unbiased estimates. In contrast, net diversification, defined by the difference in *λ* and *µ* is even less well estimated than either alone: a too large number is subtracted from a too small number. As a result, empirical results that summarize diversification using turnover are likely to be less biased than those relying on net diversification, especially in small clades and/or when the underlying extinction fraction is high. In studies of modern taxa net diversification rate differences remain a perennial favorite estimator, despite some encouragement to use other parameters for biological realism (Beaulieu and O’Meara 2016; Vasconcelos et al. 2022b). In paleontology, there is considerably more enthusiasm for using net turnover, or its reciprocal, average species lifespan (Vrba 1993).

With respect to structural bias, conditioning the likelihood on a censored tree set changes the probability space on which the model is defined. Because two-taxon “cherry trees” lack information to jointly identify distinct speciation and extinction rates, conditioning on both crown survival and *n* > 2 restricts the likelihood to trees that contain information about both parameters. We found that this conditioning scheme reduced inflation of rate estimates when small clades were censored. The analytical basis for the limited information in cherry trees is reflected in their likelihood surface.

These findings have direct implications for birth-death models that allow for heterogeneity in rates across a given tree. To estimate birth, death, or net diversification rates with even a hope of similarity to the truth one must analyze clades of at least three taxa; this holds true even if the clade of interest is inside a much larger tree. Methods that try different discrete regime paintings on the tree, like BAMM (Rabosky 2014), ClaDS (Maliet et al. 2014), and MEDUSA (Alfaro et al. 2009) could in theory be configured such that any examined regimes have the requisite amount of information. This can also be used for models that associate states with rates. For example, BiSSE could be limited to only trying distinct rates for states present in at least three tips. Difficulties come with methods that do not allow such discrete limitations. For example, with MiSSE (Vasconcelos et al. 2022a), rates are associated with hidden states that are reconstructed at tips and at nodes: there is no fixed painting of regimes, only statistical inferences about likelihoods.

Note that Bayesian methods are not an automatic fix to these issues. If a likelihood estimator is biased, weighting it by a Bayesian prior does not eliminate this bias for usual priors, though one could attempt to create various priors with biases or constraints to “correct” the bias. For both likelihood and Bayesian approaches, the best approach might be to estimate parameters as normal (subject to constraints about tree size) but then use the corrections above to get more unbiased estimates.

Likelihood estimators remain a powerful tool for inferring diversification dynamics, but their biases are most apparent in small trees or small subclades within larger trees. Recognizing and correcting these biases is essential for accurate rate estimation and for the broader application of birth–death models in comparative analyses.

## Acknowledgments

We thank James Boyko, Nathanael Walker-Hale, and Andrew Alverson for discussions and edits that have improved the content and presentation. We are also deeply indebted to the two reviewers for pointing out important issues that greatly improved the mathematical details presented here.

## Data Availability

All scripts and substantial outputs are available at https://github.com/thej022214/BeaulieuOMeara_BDBiasCode.

## A Identifiability limits in cherry trees

To explore whether cherry trees contain sufficient information to separately estimate speciation and extinction rates, we focus on the constant-rate birth-death likelihood in Eq. 3, which conditions on the survival of the crown and relies on a product over the edges as opposed to internal nodes. By reducing this expression to the special case of a two-taxon tree, we can isolate the contribution of a single observed bifurcation event and ask whether the data it contains can, in principle, identify both parameters. This simplified setting serves as a useful conceptual test case: even with a likelihood that can be evaluated, do cherry trees carry enough information to distinguish between speciation and extinction?

First, since we are examining a cherry tree, we can set *m* = 1, *t*_*i,r*_ = *T* and *t*_*i,t*_ = 0 in Eq. 3. Focusing on the product term on the left-hand side, we can simplify this expression based on what each edge of the two edges in a cherry tree contributes:

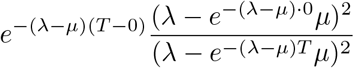

which simplifies further to,

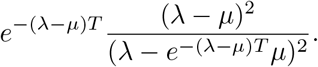

There are two such terms, one for each edge, so we square this

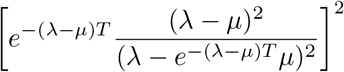

and this simplifies to,

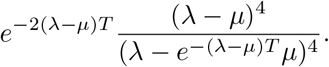

Since *m* = 1, the right hand term in Eq. 3 that includes the conditioning scheme simplifies to:

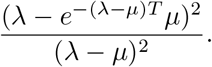

The full likelihood therefore is,

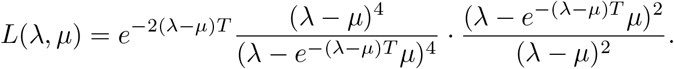

If we multiply the product term and the conditioning term together and further simplify, we obtain a final likelihood expression for a cherry tree as

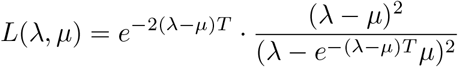

Finally, we convert this function to a log-likelihood:

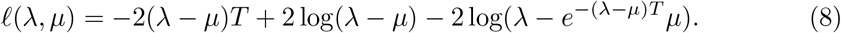

To assess identifiability, we examined the shape of the log-likelihood surface by taking the partial derivative of Eq. 8 with respect to *λ*, holding *µ* constant. The derivative is:

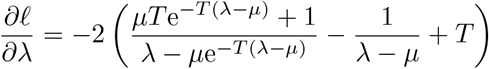

This expression describes how the likelihood changes with *λ*, and allowed us to determine whether the function has a maximum, is flat, or decreases monotonically.

We evaluated the first derivative of Eq. 8 on a grid of (*λ,µ*) values at fixed time depths, *T* =1 and *T* =10. For both depths examined, the surface of *∂𝓁/∂λ* is strictly nonzero across the parameter space, except at the singularity *λ* = *µ* (Fig. 3). As *T* increases, the magnitude of the derivative increases, indicating that the log-likelihood becomes steeper with respect to *λ*. This confirms that, for any fixed value of *µ*, the log-likelihood is a strictly decreasing function of *λ*, with no stationary point in the *λ* direction. The absence of a maximum implies that *λ* and *µ* are not jointly identifi-able from a single bifurcation in a two-taxon tree. Although a likelihood surface can be computed, it contains insufficient information to distinguish between alternative combinations of speciation and extinction rates.

**Fig. 3:**
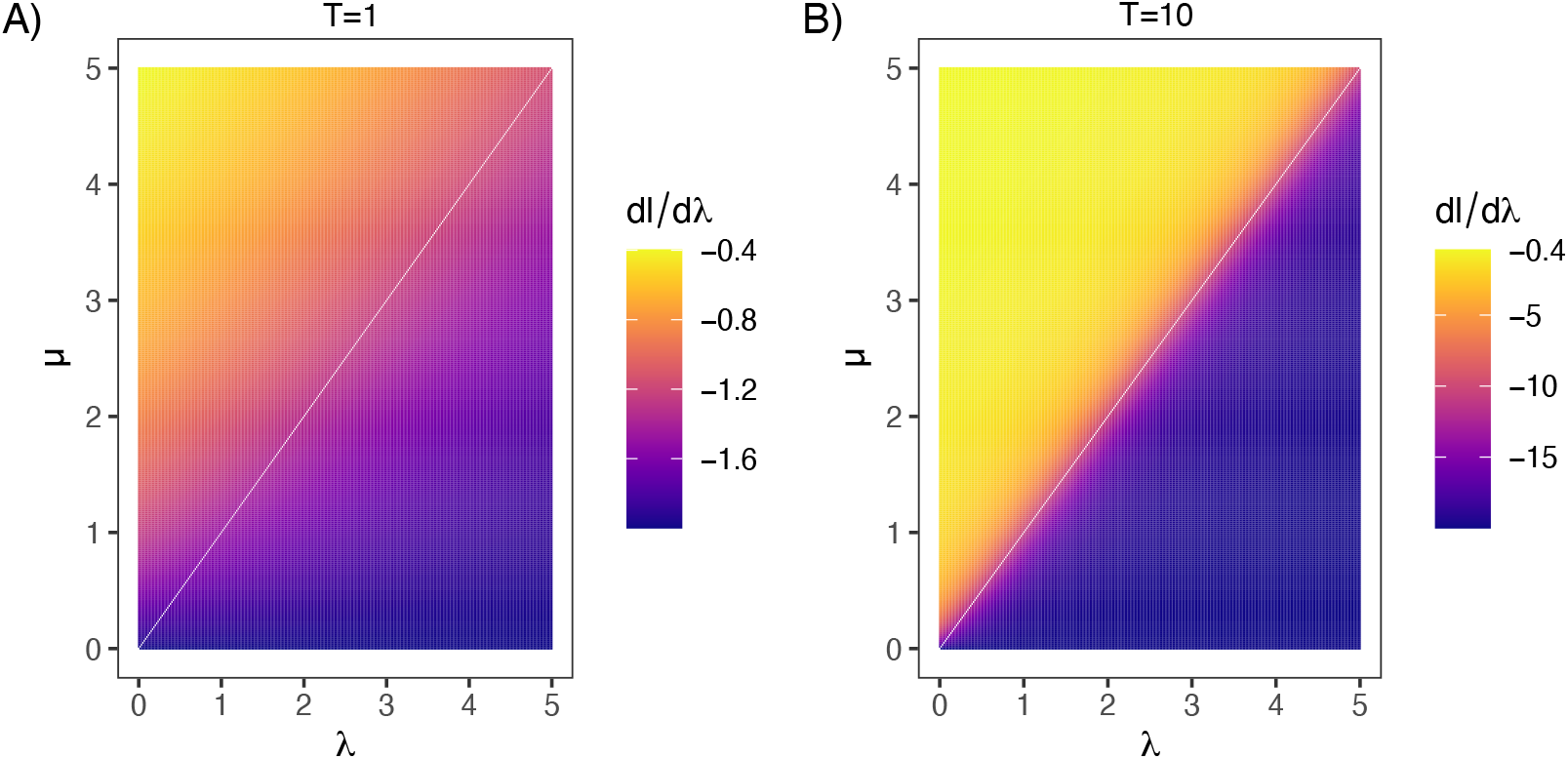
Heatmap plots of the partial derivative of the two-taxon log-likelihood function with respect to the speciation rate, *λ*, across a grid of *λ* and *µ* values. Colors represent the value of the derivative, with darker shades indicating steeper gradients. The left panel shows results at time depth *T* =1, and the right panel at *T* =10. A green contour line marks where the derivative equals zero. In both cases, the derivative is strictly negative throughout the parameter space except along the singularity at *λ*=*µ*, suggesting the likelihood estimate for *λ* must be at the lower bound, zero.

The derivative with respect to *µ* is more nuanced. It is given by:

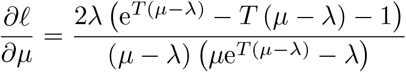

Unlike *∂𝓁/∂λ*, which is strictly nonzero across the parameter space, *∂𝓁/∂µ* exhibits regions where it approaches zero. These near-zero values occur when extinction far exceeds speciation (i.e., *µ* ≫ *λ*), particularly at deeper time depths (*T* =10)(Fig. 4). In these regions, the log-likelihood surface is relatively flat in the *µ* direction, indicating weak sensitivity to changes in *µ* and thus weak identifiability. Importantly, the derivative also flattens in the region where *λ* > *µ*, which is often more biologically plausible.

**Fig. 4:**
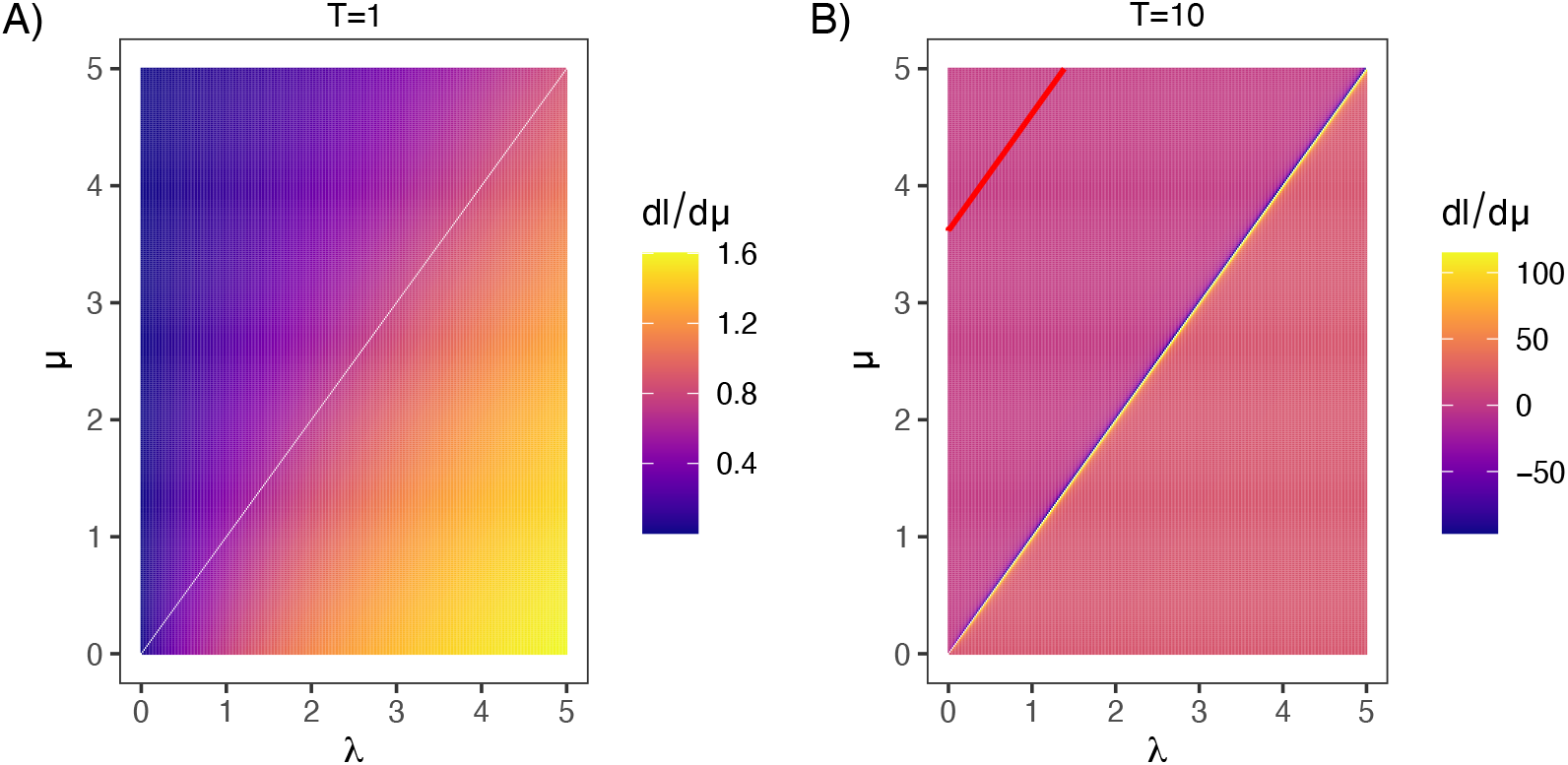
Heatmap plots of the partial derivative of the two-taxon log-likelihood function with respect to the extinction rate, *µ*, across a grid of *λ* and *µ* values. Colors indicate the magnitude of the derivative, and the dark red contour line highlights where the derivative equals zero. The left panel corresponds to a time depth of *T* =1, and the right panel to *T* =10. In contrast with the derivative with respect to *λ*, the derivative with respect to *µ* includes regions where the surface is nearly flat (i.e., derivative near zero), particularly when *µ* ≫ *λ*. These regions become more prominent for older trees, indicating that changes in *µ* have diminishing influence on the likelihood surface in those parts of parameter space. This pattern suggests that extinction rates may be weakly identifiable in some scenarios, though no true maximum is present, reinforcing the overall conclusion of non-identifiability from two-taxon trees.

Together, these results demonstrate that the likelihood function for a two-taxon tree lacks a maximum in either parameter dimension, confirming that speciation and extinction rates cannot be jointly identified from a single bifurcation.

## B Symbolic regression grammar

Symbolic regression was implemented using the `gramEvol` R package (Noorian et al. 2016). The analysis used the following context-free grammar, which defines the space of candidate expressions evaluated. For 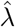 in the birth-death case, we used the following set up:

~~~
grammarDef <-CreateGrammar(list(
    xpr = grule(op(expr, expr), func(expr), var),
    func = grule(log1p),
    op = grule(‘+’, ‘-’, ‘*’, ‘/’),
    var = grule(lambda_fit, eps_fit, ntips,
                Ntips_over_Ntips_minus1,
                Ntips_minus1_over_Ntips_minus2,
                const),
    const = grule(0.5, 1, 2, 3)
))
~~~

The grammar permits basic arithmetic operators, a single nonlinear transformation (log(1 + *x*)), and a restricted set of variables and constants. Variables include the likelihood estimate of the speciation rate 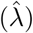, the estimated extinction fraction 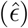, the number of extant taxa (*n*), and two dimensionless sample-size correction terms, *n/*(*n* − 1) and (*n* − 1)*/*(*n* − 2). Constants were restricted to a small discrete set to limit redundancy. All expressions considered in the analysis were required to take a multiplicative correction form, 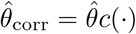, ensuring dimensional consistency.

